# A deep learning-based multiscale integration of spatial omics with tumor morphology

**DOI:** 10.1101/2024.07.22.604083

**Authors:** Benoît Schmauch, Loïc Herpin, Antoine Olivier, Thomas Duboudin, Rémy Dubois, Lucie Gillet, Jean-Baptiste Schiratti, Valentina Di Proietto, Delphine Le Corre, Alexandre Bourgoin, Pr. Julien Taïeb, Pr. Jean-François Emile, Pr. Wolf H. Fridman, Elodie Pronier, Pr. Laurent-Puig, Eric Y. Durand

## Abstract

Spatial Transcriptomics (spTx) offers unprecedented insights into the spatial arrangement of the tumor microenvironment, tumor initiation/progression and identification of new therapeutic target candidates. However, spTx remains complex and unlikely to be routinely used in the near future. Hematoxylin and eosin (H&E) stained histological slides, on the other hand, are routinely generated for a large fraction of cancer patients. Here, we present a novel deep learning-based approach for multiscale integration of spTx with tumor morphology (MISO). We trained MISO to predict spTx from H&E on a new unpublished dataset of 72 10X Genomics Visium samples, and derived a novel estimate of the upper bound on the achievable performance. We demonstrate that MISO enables near single-cell-resolution, spatially-resolved gene expression prediction from H&E. In addition, MISO provides an effective patient representation framework that enables downstream predictive tasks such as molecular phenotyping or MSI prediction.

## Introduction

Classical transcriptomics methods like microarray and bulk RNA sequencing have been crucial in cancer research^1^, aiding in understanding tumorigenesis^2–4^, tumor heterogeneity^5^, and developing treatments. However, these techniques merge information from diverse cell types and structures within a tissue sample, masking signals from rare cell populations and the spatial dynamics around tumors. The functionality of a cell type is influenced by signals from neighboring cells^6,7^, highlighting the importance of spatial context in immune cell response and tumor microenvironment arrangement for understanding tumorigenesis^8^, progression^9^, and patient prognosis^10^.

Advanced technologies such as 10X Genomics Visium and 10X Xenium offer higher resolution spatial transcriptomics, but are costly and not widely used in clinical settings. In contrast, haematoxylin and eosin (H&E)-stained slides remain a diagnostic standard in cancer care. Recent advancements in Artificial Intelligence (AI) have shown that high-resolution H&E images can predict molecular features like mutations^11^, molecular phenotypes^12,13^, bulk transcriptomic expression^14^ and patient outcomes^15^.

The increased availability of spTx datasets has fueled the development of models that predict spatially resolved transcriptomic features from H&E slides. Current spTx technologies like Visium measure gene expression in “spots” (55µm in diameter for Visium). On H&E whole-slide images (WSI), small patches called tiles (typically of size 112µm x 112µm) are generally used as inputs to models, and can easily be matched with spots. This framework promises to enhance spatial transcriptomics data, potentially increasing resolution to near single-cell levels and applying to vast collections of H&E slides at minimal additional cost. Prior methods have attempted to predict gene expression at the tile level, without encoding the local spatial arrangements between tiles^16,17^. More recent methods attempt to capture the spatial organization of WSI by leveraging neighborhood information^18–21^.

However, most current approaches rely on either small or low-resolution datasets of spatial transcriptomics data^17,22–24^ limiting their robustness on external validation cohorts. Altough recently, Jaume *et al*. introduced HEST-1K^25^, a large dataset of 1108 samples, to address this limitation, this dataset is heterogeneous in terms of data acquisition technology, resolution, species (most samples coming from mice), and therapeutic domain. Specifically in colorectal cancer, HEST-1K contains 19 samples from less than 10 patients.

Here, we introduce MISO (Multiscale Integration of Spatial Omics), a novel and unified deep learning framework to integrate spatial knowledge at multiple scales, from the cell to the patient. We leveraged a dataset (PETACC8-Visium) of 72 Visium samples of distinct patients with colorectal cancer, to our knowledge the largest curated dataset in this domain, to train and robustly assess the out-of-domain performance of our approach.

MISO relies on a local self-attention mechanism to model the spatial organization of the tissue, allowing to infer close to single-cell resolution gene expression on datasets where only H&E slides are available and enriching the representation of patient data (Figure 1). In order to take into account the patient-to-patient variability, we used a loss function specifically designed to tackle this effect. Relying on a robust statistical framework, we derived theoretical upper bounds to the performance of models predicting spatial gene expression. We trained models not only to predict gene expression at the level of spots, but also to learn a finer-grained resolution of spatial omics data, effectively approaching the resolution of spatial single-cell sequencing. Once trained MISO only requires H&E images to infer predictions on new samples (out-of-domain prediction), while most alternative super-resolution approaches also require Visium data to perform the same kind of inference (in-domain prediction). Finally, models trained on the PETACC8-Visium cohorts were used to infer a high-level representation of slides of colon adenocarcinoma from The Cancer Genome Atlas (TCGA-COAD). Even though this representation was trained on only 48 samples, it proved superior to the conventional representation of histology slides for several downstream predictive tasks.

**Figure 1.**
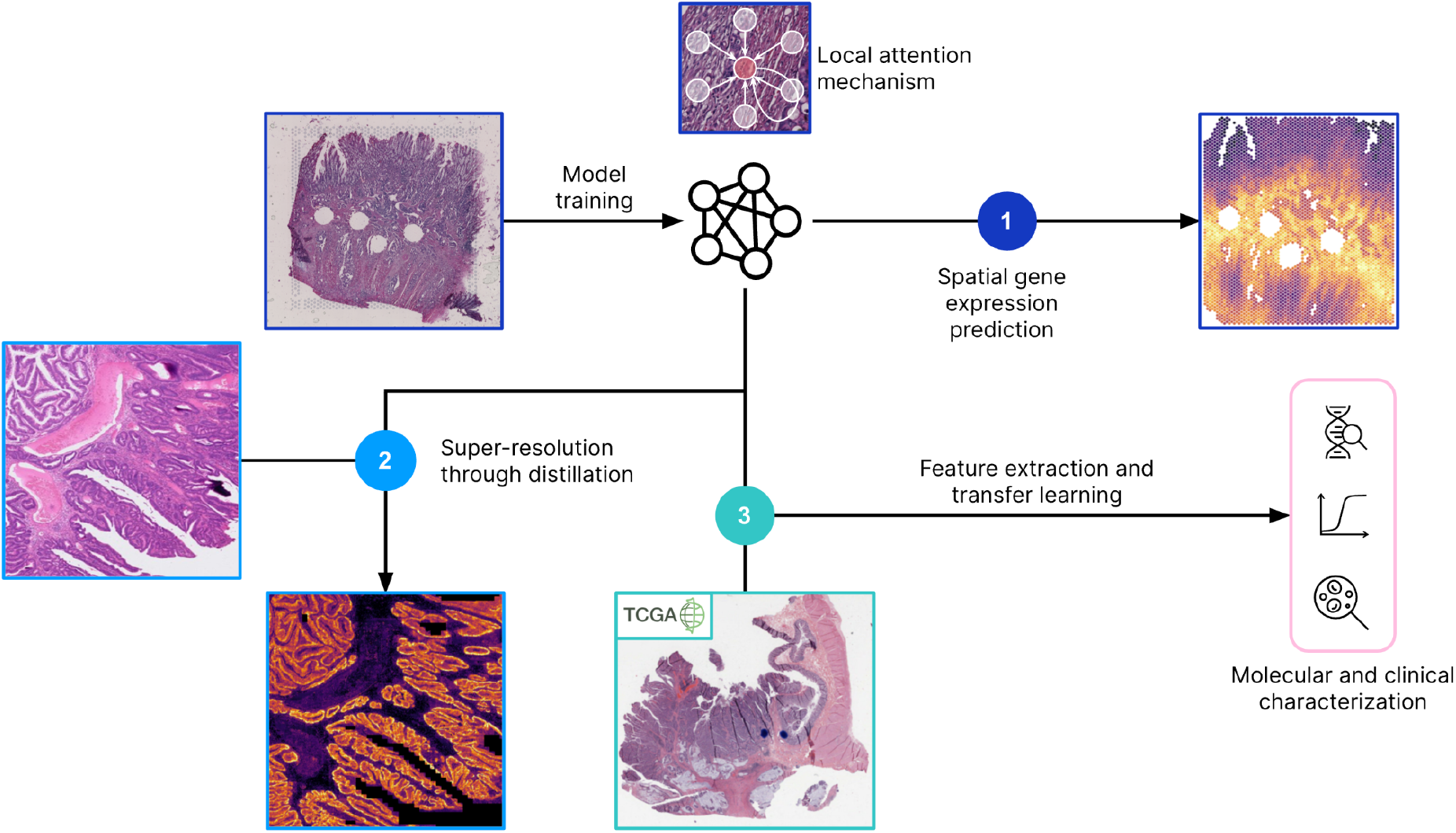
Graphical summary. (1) Models with the local attention MIL architecture were trained to predict the spatial expression of genes in a cohort of 48 samples with 10X Visium sequencing. (2) Those models were used as “teachers” in a distillation setting: they were used to generate pseudo-labels in the same cohort, and a lighter architecture was trained to predict these pseudo-labels with weak supervision, allowing to generate predictions with a finer-grained resolution. These high resolution predictions were validated against cell type annotation and 10X Xenium sequencing. (3) The models trained to predict spatial gene expression were used to extract features in TCGA cohorts. The extracted features were compared to the conventional features trained in a self-supervised manner using H&E images only in several downstream tasks.

## Results

### MISO predicts spatial gene expression from histology images

We propose MISO, a deep learning-based model to predict spTx from histology. MISO was trained on a dataset of 48 samples of colorectal cancer patients, sequenced with 10X Visium technology. For each slide, tile images centered on the location of Visium spots (up to 5,000 per slide) were selected. Features were extracted from tiles with a Vision Transformer model (iBOTViTBase), pretrained with self-supervised learning on TCGA-COAD (see Methods).

We compared two approaches: similar to HE2RNA^14^ and STNet^16^, a baseline MLP without any spatial context, which takes as input the features of a tile and predicts the expression of genes in the corresponding spot, and a transformer-based architecture that integrates spatial interactions by computing self-attention terms between every pair of inputs (see the Methods for details). Because it is computed on every input, classical self-attention has two main limitations: it does not discriminate between long- and short-range interactions and it has a high computational burden because of its quadratic complexity. The Local Attention Multiple Instance Learning (LAMIL) model^26^ overcomes these limitations by restricting the computation of cross-attention scores to neighboring tiles only. MISO is an adaptation of this architecture to the task of predicting gene expressions of each tile. This significantly reduces the computational burden compared to generic transformers, while keeping relevant information, as we expect gene expression in one spot to be influenced by neighboring spots due to cell-cell interactions. A key hyperparameter of this method is the number *k* of neighbors used in the self-attention computation, higher values covering larger areas while increasing the computational cost.

Importantly, no consensus has been reached regarding the processing and modeling of spTx data. In previous works, pre-processing methods vary from simply using raw data to log-CPM normalization, SCTransform^27^ or min-max scaling. However, to the best of our knowledge, all those methods fail to address inter-patient variability, thus classical regression models trained with Mean Squared Error may be confounded by this patient effect. Here, we propose a different approach by training our models with a loss based on cosine similarity to maximize for each gene and each sample the correlation between predictions and raw counts. This means that each slide from the training set had to be processed at once in the same batch. As shown in the Methods section, models trained with this loss function were indeed more robust.

For each architecture, the models were trained on 25 random train/test splits of the data. To evaluate performances, we computed average Spearman and Pearson correlation per gene/slide pair on the test set.

At first, to evaluate the performance of our method on a small curated list of genes, we restricted our analysis to the set of 100 genes with the highest spatial autocorrelation. For this, we computed the Moran’s I of each gene separately on every slide, and we used the geometric mean over samples to aggregate the results over the dataset.

For MISO, we varied the number *k* of neighbors used in self-attention computation from the list (0, 6, 18, 36, 60, 90 and 126), corresponding to a growing hexagon around the spot. With *k* = 0, MISO (average Spearman correlation = 0.347, st = 0.052, average Pearson correlation = 0.328, std = 0.049) and the baseline architecture (average Spearman = 0.343, st = 0.053, average Pearson = 0.328, std = 0.045) obtained consistent performances. The performance of MISO increased first quickly with the number of neighbors, up to a plateau between *k* = 18 (average Spearman = 0.366, std = 0.055, average Pearson = 0.345, std = 0.047) and *k* = 126 (average Spearman = 0.370, std = 0.056, average Pearson = 0.347, std = 0.048), demonstrating the relevance of neighborhood interaction for the determination of local gene expression (comparison between the best architecture and the baseline: p = 2 × 10^−3^).

Next, we extended the list to the 5,000 genes with the highest spatial autocorrelation. The same experiment of varying the value of *k* gave consistent results with the previous one, the peak of performance being reached here with *k* = 36, with an average Spearman correlation of 0.219 (std = 0.041) (average Pearson = 0.217, std = 0.034).

For a given gene and a given sample, the performance in predicting the gene expression was limited by statistical noise in the labels. We derived a novel estimate of the Pearson correlation that would be achieved by a perfect oracle having access to all deterministic variation factors, that we denote *R*_*max*_ (see Methods). This value sets an upper bound on the Pearson correlation *R* achieved by a model (Figure 2). To investigate the biological processes best captured by the model, we ranked genes based on the ratio *R* / *R*_*max*_, and ran an enrichment analysis of genes with the highest ratio (Figure 2, see Methods). The most enriched pathway was that of genes regulated by MYC, a well-known proto-oncogene (including MYC itself). Previous work has found that MYC was up-regulated in all stages of colorectal cancer, where it was the main source of metabolic reprogramming^28^. Other enriched pathways consistently covered metabolic processes, but also mechanisms involved in cancer development (resistance to programmed cell death for instance), and normal functions of colon mucosa (epithelium development). Among those pathways, some are known to be influenced by their spatial localization. Metabolic processes for example are affected by the hypoxic and necrotic level of the tumor which are both often higher in the tumor core than in its margin.

**Figure 2.**
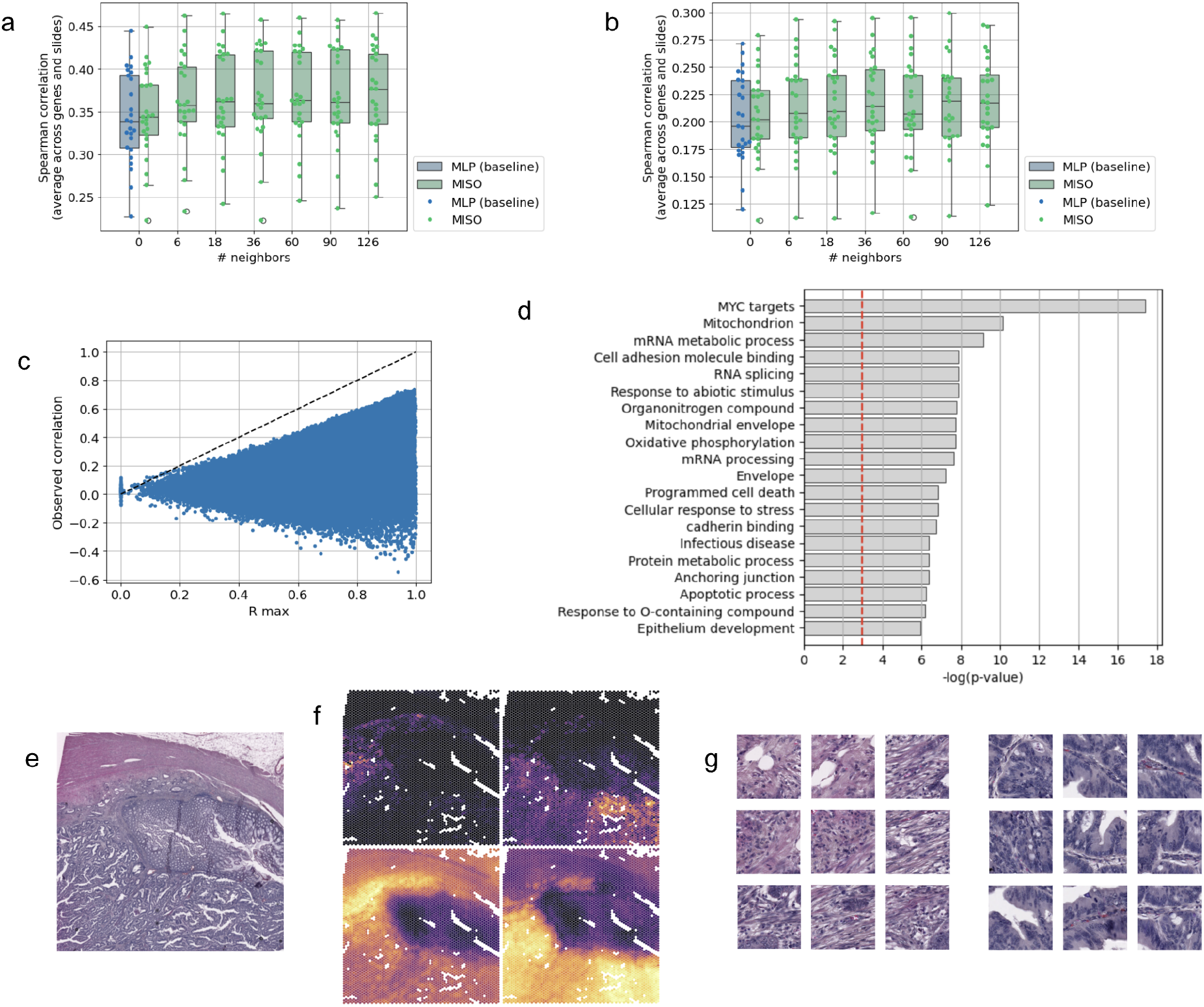
Spatial gene expression prediction. **a**. Boxplot of the performances obtained for the prediction of the 100 genes with highest Moran’s I in repeated train/test splits of the 48 training samples depending on the architecture and the number of nearest neighbors in the case of Local Attention MIL. **b**. Same, for the prediction of the 5,000 genes with the highest Moran’s I. **c**. Pearson correlation achieved by the 5,000-gene model for every gene/slide pair, against the theoretical upper bound derived in Methods - Modeling of Spatial Transcriptomic data. The dotted line indicates the performances that would be achieved by a perfect oracle. **d**. Gene set enrichment analysis on the 118 genes best predicted with regard to this upper bound. **e**. Capture area on a sample from the external test set. **f**. Ground truth average expression of *COL1A1* and *COL1A2* (top left) and *KRT8* and *KRT18* (top right) and predictions of the model for the same genes (bottom) on the same sample. **g**. Tiles with highest predicted expression of *COL1A1* and *COL1A2* (left), and *KRT8* and *KRT18* (right) on the same sample.

A dataset of 24 additional samples, made available only after training the models, was used as a validation cohort. During inference, the 25 models trained on each test splits were applied on new samples, and their predictions were averaged. Consistently with cross-validation results, MISO outperformed the baseline both for the prediction of the 100 and 5,000 most varying genes (p-value < 1 × 10^−10^ for both lists) (Table 1). These results validate the robustness of our approach in an unseen dataset.

**Table 1:** performances of the models in cross-validation and external validation. For internal validation, the average and standard deviation of each metric on the 25 test splits are reported. For external validation, the correlation and the 95% confidence interval, obtained by bootstrapping, are reported. On Xenium samples, we report the average and standard deviation of the correlation over genes.

| # genes | Setting | Model | Spearman | Pearson |
| --- | --- | --- | --- | --- |
| 100 | Internal validation<br>(N = 48) | MLP (baseline) | 0.343 (0.053) | 0.328 (0.049) |
| | | MISO ($k = 126$) | 0.370 (0.056) | 0.347 (0.048) |
|  | External validation<br>(N = 24) | MLP (baseline) | 0.430 [0.421 - 0.438] | 0.431 [0.423 - 0.439] |
| | | MISO ($k = 126$) | 0.491 [0.482 - 0.500] | 0.468 [0.460 - 0.477] |
| 5000 | Internal validation<br>(N = 48) | MLP (baseline) | 0.203 (0.039) | 0.201 (0.037) |
| | | MISO ($k = 36$) | 0.219 (0.041) | 0.217 (0.034) |
|  | External validation<br>(N = 24) | MLP (baseline) | 0.256 [0.255 - 0.257] | 0.268 [0.267 - 0.269] |
| | | MISO ($k = 36$) | 0.284 [0.283 - 0.285] | 0.279 [0.278 - 0.280] |
| 146 | Xenium colorectal<br>(8μm x 8μm<br>patches) | iStar | 0.169 (0.120) | 0.142 (0.103) |
|  |  | MISO (distil.) | 0.220 (0.209) | 0.186 (0.173) |
| 105 | Xenium breast<br>(6μm x 6μm<br>patches) | iStar | 0.139 (0.144) | 0.136 (0.143) |
|  |  | MISO (distil.) | 0.151 (0.139) | 0.147 (0.143) |
| | Xenium breast<br>(12μm x 12μm<br>patches) | iStar | 0.134 (0.132) | 0.136 (0.140) |
|  |  | MISO (distil.) | 0.222 (0.204) | 0.218 (0.217) |

**Table 2:** Comparison of performances of models trained on top of the conventional iBOTViTBase representation or of the MISO-based representation, on subsampled datasets of 200 patients and on the entire TCGA cohorts.

| Cohort | Task (metric) | Features | Population size |  |
| --- | --- | --- | --- | --- |
|  |  |  | 200 patients | All patients |
| TCGA-COAD | Hallmarks (Spearman) | iBOTViTBase | 0.148 (0.017) | 0.269 (0.008) |
|  |  | MISO | 0.209 (0.020) | 0.279 (0.009) |
|  | CD3 - CD19 - CD20 (Spearman) | iBOTViTBase | 0.295 (0.054) | 0.419 (0.026) |
|  |  | MISO | 0.363 (0.066) | 0.451 (0.023) |
|  | Survival (C-index) | iBOTViTBase | 0.629 (0.067) | 0.659 (0.014) |
|  |  | MISO | 0.633 (0.064) | 0.661 (0.021) |
|  | MSI (AUC) | iBOTViTBase | 0.907 (0.029) | 0.941 (0.007) |
|  |  | MISO | 0.901 (0.029) | 0.938 (0.005) |
|  | CMS (AUC - macro-average) | iBOTViTBase | 0.790 (0.020) | 0.819 (0.008) |
|  |  | MISO | 0.800 (0.023) | 0.824 (0.006) |
| TCGA-STAD | MSI (AUC) | iBOTViTBase | 0.819 (0.045) | 0.866 (0.013) |
|  |  | MISO | 0.836 (0.039) | 0.876 (0.009) |
|  | Survival (C-index) | iBOTViTBase | 0.590 (0.032) | 0.595 (0.017) |
|  |  | MISO | 0.588 (0.038) | 0.603 (0.014) |
| TCGA-BRCA | Hallmarks (Spearman) | iBOTViTBase | 0.136 (0.024) | 0.360 (0.004) |
|  |  | MISO | 0.222 (0.028) | 0.366 (0.005) |
|  | CD3 - CD19 - CD20 (Spearman) | iBOTViTBase | 0.347 (0.074) | 0.564 (0.005) |
|  |  | MISO | 0.423 (0.066) | 0.570 (0.007) |
|  | Survival (C-index) | iBOTViTBase | 0.591 (0.107) | 0.659 (0.017) |
|  |  | MISO | 0.593 (0.112) | 0.649 (0.016) |
|  | HRD (AUC) | iBOTViTBase | 0.700 (0.071) | 0.791 (0.007) |
|  |  | MISO | 0.699 (0.072) | 0.792 (0.008) |
|  | BRCA1/BRCA2 (AUC) | iBOTViTBase | 0.549 (0.134) | 0.618 (0.020) |
|  |  | MISO | 0.550 (0.124) | 0.623 (0.026) |
|  | Subtype (AUC - macro-average) | iBOTViTBase | 0.753 (0.038) | 0.842 (0.004) |
|  |  | MISO | 0.766 (0.033) | 0.841 (0.004) |

In summary, MISO proved to be a reliable model to predict spatial gene expression in colorectal cancer patients.

### From spot resolution to spatial super-resolution

SpTx technologies (like Visium) offer unprecedented insights about the relation between gene expression and tissue morphology. However, they do not reach spatial single-cell resolution. Recently, several technologies of single-cell-level spatial transcriptomics have been developed, such as 10X Genomics Xenium or nanoString’s CosMx, but they remain particularly expensive, and restricted in the number of transcripts that can be measured simultaneously.

Previous attempts have been made to bridge the gap between these two levels of resolution, such as BayeSpace^29^, XFuse^30^ or iStar^31^. Yet, they still carry some limitations as they are trained on very small datasets, and their evaluation relies on in-sample performance. For instance, iStar’s design requires “low-resolution” sequencing data matched to an H&E slide (or to a consecutive slice) to be able to infer a super-resolved gene expression.

Here, we built upon our supervised local-attention transformer model to tackle this task of robustly inferring super-resolved expression maps from histology slides alone, as opposed to methods requiring spTx data in inference. Following previous work^14,31^, we leveraged weakly supervised learning to increase the spatial resolution of available sequencing, by further dividing tiles into 196 (14×14) patches of size 8μm x 8μm (see Methods).

In practice, the augmentation of the resolution is computationally expensive. To circumvent this, we employed a knowledge distillation approach^32^ (see Methods). Spot-level predictions of the previously trained supervised model (“teacher” model) were used as pseudo-labels to train a lighter architecture (“student” model).

To assess the spatial accuracy of our approach, we first evaluated qualitatively the performance on an in-house dataset of tiles from TCGA-COAD samples for which nuclei were annotated by pathologists (see Methods). We measured the association between these nuclei annotations and our higher-resolved expression maps. A patch was considered positive for a given nucleus type if it overlapped with its segmentation mask. As expected, *KRT8* and *KRT18* (keratin markers) expression was higher in patches positive for epithelial cells, both healthy (p-value < 10^−10^, one-tailed t-test) and tumoral ones (p-value < 10^−10^), while *COL1A1* and *COL1A2*, markers of collagen, were more expressed in patches positive for fibroblasts (p-value < 10^−10^). We also investigated the predicted expression of genes coding for T-cell and B-cell receptors. Despite the fact that those genes had very low expression levels in the training set and that the teacher model achieved low performance, we found that the student models predicted significantly higher expression of *CD3* and *CD19* genes in lymphocytes patches than in any other cell type (p-value < 10^−10^) (Figure 3).

**Figure 3.**
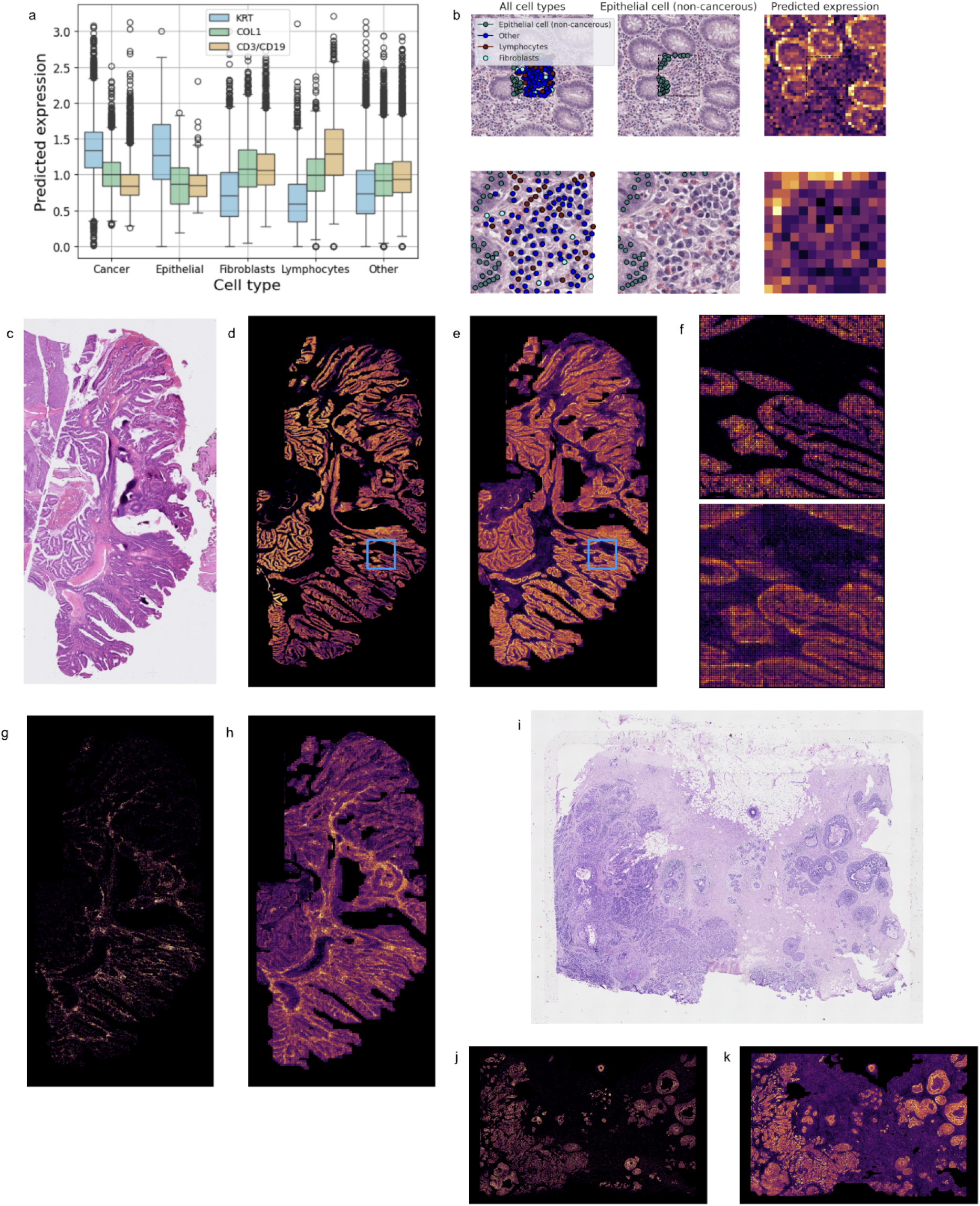
Validation of the super-resolved expression maps. **a**. Boxplot of the predicted expression of marker genes in various cell types in TCGA-COAD annotated tiles. *COL1*: average expression of *COL1A1* and *COL1A2*. KRT: average expression of *KRT8* and *KRT18. CD3*/*CD19*: average expression of *CD3D, CD3E, CD3G, CD247* and *CD19*. **b**. Example tile annotated with non-tumor epithelial cells and other cell types with (top) and without (bottom) its context, and predictions of the model for the *KRT8* and *KRT18* genes on patches of size 8µm x 8µm (left columns). **c**. H&E sample of colorectal cancer. **d**. Ground truth expression of *EPCAM* obtained with the 10X Xenium technology. For comparison with the model’s predictions, transcripts were summed on patches of size 8µm x 8µm. **e**. Prediction of the expression of *EPCAM* by the MISO super-resolution approach. **f**. Ground truth (top) and predicted (bottom) expression of *EPCAM* in the area indicated by the blue squares in b. and c. **g**. Ground truth expression of T cell and B cell markers (average expression of *CD3D, CD3E, CD247, CD19 and MS4A1*). **h**. Predicted expression of T cell and B cell markers (average expression of *CD3D, CD3E, CD3G, CD247* and *CD19*). **i**. H&E sample of breast carcinoma. **j**. Ground truth expression of *CDH1* obtained with the 10X Xenium technology, as in d. **k**. Prediction of the expression of *CDH1* by the MISO super-resolution approach.

We further validated quantitatively our super-resolution approach on Xenium samples, that provide a ground truth at single-cell resolution. We used two samples: one of colorectal cancer and one of breast cancer (Figure 3). 480 genes were sequenced in each sample, among which 146 were also in our 5,000 gene list for the colon sample and 105 for the breast sample. We trained models to predict the expression of the common genes. We benchmarked our method on this task against the iStar algorithm^31^. For the comparison, we retrained iStar in the same configuration as MISO, on the same 146 (resp. 105) genes. To compare the predictions generated by our models to the ground truth, we first aligned the Xenium sample to the associated H&E image. Then, we divided the WSI into small patches and summed the detected transcripts over each one.

On the colon adenocarcinoma sample, with 8µm x 8µm patches, the model achieved an average Spearman correlation of 0.220, and a Pearson correlation of 0.186 (Table 1), that can be compared to the performances of the teacher model to predict the same list at Visium resolution: resp. 0.238 and 0.235. In particular, Spearman correlations above 0.50 were achieved for 21 genes (Table S3). On the same sample, iStar achieved overall lower performances than MISO (Table 1, Fig. S3, p-value = 2 × 10^−7^), reaching Spearman correlation above 0.5 for only a single gene: ERBB3 (0.501 for iStar vs 0.672 for MISO).

Then, we applied the MISO model predicting the expressions of T cell and B cell markers. Again, despite its low spot-level performance in internal validation, this model reached significant performances at the patch level, with up to 0.208 in Spearman correlation and 0.263 in Pearson correlation for the prediction of *CD3E* in the Xenium sample. In a second step, we summed the expressions (ground truth and predicted) of all available marker genes of T cells, B cells or both. The summed prediction of the model for T cell markers achieved Spearman and Pearson correlation of 0.237 and 0.296 respectively, while the performance for B cells was more moderate (0.100 and 0.198 resp.). The prediction of summed B cell and T cell markers reached Spearman and Pearson correlations of 0.259 and 0.360 resp. (Table S3). These results highlight the fact that, while trained to predict very noisy labels, our approach allowed us to recover a biological signal.

For the Breast Carcinoma sample, the highest resolution level of the associated H&E image was 0.36 micron per pixel. We applied our models (trained at 0.5 micron per pixel) at this resolution, meaning that the smallest patches we considered were slightly smaller than 6µm x 6µm. It should be noted that both the resolution and the organ differed from the training data. We measured an average Spearman and Pearson correlation of respectively 0.151 and 0.147 (Table 1), against 0.253 and 0.249 at Visium resolution in internal validation. Once again, the expression of *EPCAM* was well predicted, with a Spearman correlation of 0.501. At this fine-grained resolution, this level of performance was not reached for any other gene, but the model achieved Spearman correlations above 0.4 for seven additional genes (Supp. Table 2). With 12µm x 12µm patches, average Spearman and Pearson correlation were respectively 0.222 and 0.218 (Table 1), and Spearman correlation above 0.5 was achieved for nine genes (Supp. Table 2). The prediction of B cell and T cell marker genes gave results very consistent with those obtained on the colorectal cancer case (Tables S3 and S4).

On the same sample, iStar reached an average Spearman correlation of 0.139 for the prediction of the same genes with 6µm x 6µm patches (without any gene above 0.5 in spearman correlation), significantly below the performance of MISO (p-value = 0.001), and the difference between both approaches increased with 12µm x 12µm patches (p-value = 7 × 10^−16^), Overall, these results validate the relevance of our approach to extract high-resolution predictions in a fully out-of-domain setting, using only H&E-stained slides.

### Spatial omics-based prediction as enriched representation of histology slides

Finally, we investigated the representation learnt by the MISO model trained to predict the expression of the 5,000 most spatially variable genes. We removed the last layer and used the output of the transformer module to compute 1024-dimensional feature vectors from tile images. By design of our model and compared to the traditional features used in computational pathology, this representation incorporates information not only about the tile itself, but also about its immediate surroundings as well as local gene expression. We compared this spTx-powered representation to the feature vectors directly extracted with iBOTViTBase-COAD for several classical slide-level downstream tasks (bulk gene expression prediction, molecular subtyping, prediction of genetic alteration and survival analysis, see Methods) in TCGA-COAD (Colon Adenocarcinoma), TCGA-STAD (gastric cancer) and TCGA-BRCA (Breast Cancer). In the latter cases, we probed the out-of-domain (OOD) robustness of the method, as MISO was trained only on colorectal cancer patient data.

For each cohort and each task, we trained Chowder^33^, a multiple-instance learning model, on top of each representation. A learning curve was built by repeating the experiment for different subsampled datasets of varying size. Robust performance estimates were obtained by repeated cross-validations (see Methods).

The MISO representation outperformed the iBOTViTBase representation for bulk gene expression prediction tasks, for each considered list of genes and each cohort. The learning curve showed that the MISO representation was particularly efficient in smaller datasets (Fig. 4). For molecular subtyping in TCGA-COAD (prediction of consensus molecular subtypes - CMS) and TCGA-BRCA, the MISO representation also showed improved performance with respect to iBOTViTBase, especially when the training dataset was small. For instance, for CMS prediction in TCGA-COAD, with n = 100, MISO reached an AUC (macro-average) of 0.743, against 0.721 for iBOT (p-value = 0.028, corrected resampled T-test). On TCGA-BRCA, with n = 200, AUC(MISO) = 0.766, AUC(iBOT) = 0.753 (p-value = 0.023).

**Figure 4.**
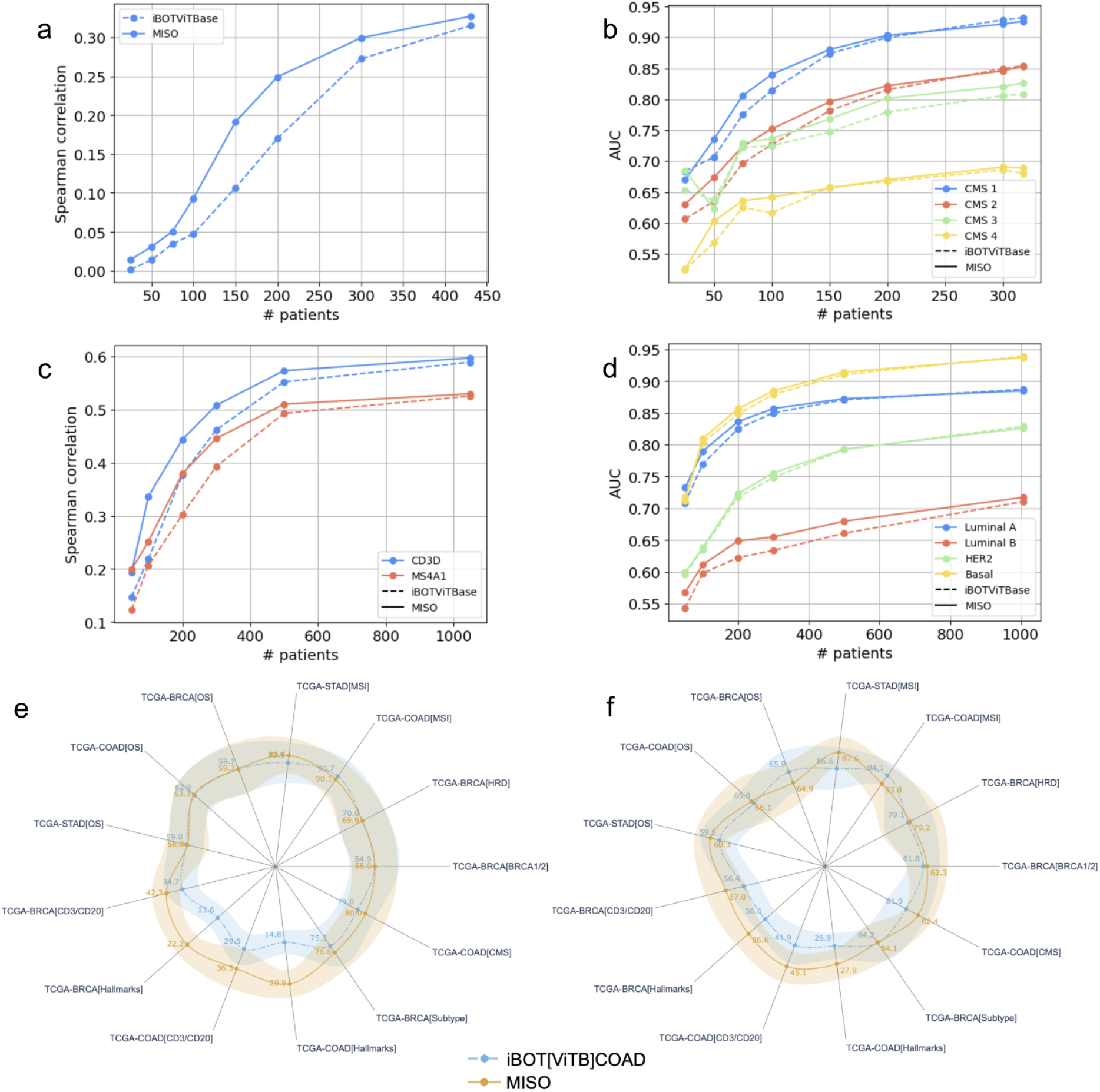
Transfer learning performances. **a**. Average Spearman correlation obtained with MISO or iBOTVit-Base representation for the prediction of the expression of T cell immunity-related genes, in TCGA-COAD, in repeated cross-validation experiments with varying dataset size. Average was computed over 20 repeats of 5-fold cross-validation. **b**. Average AUC obtained to predict each consensus molecular subtypes in TCGA-COAD. **c**. Average Spearman correlation obtained for the prediction of the expression of CD3D and MS4A1 in TCGA-BRCA. **d**. Average AUC obtained to predict each breast cancer molecular subtype in TCGA-BRCA. **e**. Comparison of the performances obtained with MISO or iBOTVit-Base representation in various transfer learning tasks, with cross-validation datasets restricted to 200 patients. The lines indicate the average performance and the shaded area the standard deviation over repeated cross-validations. **f**. Same, with all patients from TCGA-COAD, TCGA-STAD and TCGA-BRCA used for cross-validation.

For the prediction of molecular alterations (MSI, HRD, BRCA1/BRCA2 mutation), both representations offered similar performances and differences were most of the time not significant, although the MISO representation was slightly better for predicting MSI in stomach adenocarcinoma. For the prediction of overall survival, a challenging task where label noise tends to dominate performances, both representations also showed comparable results.

Overall, even if the difference was not significant for every single task, the combined p-value of all considered tasks was significantly in favor of MISO, particularly in situations where the number of training samples was limited (with n = 200 patients, p-value = 2 × 10^−15^, Stouffer test), but not only (with the whole population, p-value = 0.037).

Altogether, our results suggest that, from a given feature extractor, learning to predict Spatial Transcriptomic data, even on a relatively small dataset with only 48 patients, could help increase the performances of digital pathology models for several biologically and clinically relevant tasks.

## Discussion

We introduced MISO, a multiscale deep learning approach to explore relations between tissue morphology and local changes in gene expression. The development of our model leveraged a new dataset of 48 training samples and 24 test samples, all coming from distinct patients. This is to our knowledge the largest dataset of Visium samples from colorectal cancer patients, and one of the largest Visium datasets in oncology.

MISO was able to predict the spatial expression of a large set of genes, and one of the main limiting factors for performance was the quality of the labels obtained through Visium technology, as it is prone to shot noise due to the finite sequencing depth.

Through distillation technique, MISO was able to further refine the spatial resolution of the sequencing, effectively approaching the scale of spatial single-cell RNAseq, as validated qualitatively and quantitatively against cell annotations performed by trained pathologists and against 10X Xenium sequencing data. Compared to previous works that investigated super-resolution of spatial transcriptomic, our approach benefitted from a rich training dataset, allowing for its direct application on new cohorts where only H&E-stained histology slides are available. We emphasize the importance of this point, as many of the super-resolution approaches developed in the past rely on training the method de novo on a sample for which Visium sequencing is available.

As a direct application, we showed that MISO provided a powerful representation of cancer patient data, improving or matching the downstream performance on several histology-based machine learning tasks. In particular, it significantly enhanced the prediction of bulk gene expression, which is still a clinically relevant task as non-spatial RNAseq is involved in a variety of companion diagnostics (for instance PD-L1 expression). The representation learnt by MISO also proved powerful for downstream tasks further away from its training domain, not only in colorectal cancer, but also in breast cancer and gastric cancer.

Yet, the number of individual training samples used in the present work remains small with regards to the amount of data usually required for the training of deep learning models. Thus, we expect that there is still a large margin for improvement in the various tasks investigated here, which will be filled by increasing the number of samples.

The promising performance of our approach should encourage future works to explore the potential of spatially-resolved omics data on large datasets. By leveraging the highly-resolved data on already existing large H&E datasets (e.g. TCGA), one could explore scientific avenues unlocked by the availability of spatial transcriptomics. Such examples could include finding context-specific drug-target matches such as netrin-1 / NP137 monoclonal antibody identified by Lengrand et al. (2023)^34^ and Cassier et al. (2023)^35^, identifying small and localized immune niches (Madissoon et al. 2023)^36^, or investigating cell-cell communication “language” and its consequences on the cell state polarization and their functional states (Lyubetskaya et al. 2022)^37^.

## Data Availability

The PETACC8-Visium dataset is available from the Fédération Francophone de Cancérologie Digestive (FFCD) which is responsible for granting use of rights and accepting dissemination. The data were used with permission for the current study and are not publicly available. Such dissemination must also be compliant with data privacy laws and framework (including patient consent and information). Any reuse of the data must be approved by the FFCD and the ethics committee.

The Xenium samples are publicly available and can be downloaded from the 10X Genomics platform https://www.10xgenomics.com/datasets.

TCGA data is publicly available and can be downloaded from the GDC data portal https://portal.gdc.cancer.gov/.

## Acknowledgments

The results published here are based on a subset of a cohort from a clinical trial (PETACC-8) sponsored by the Fédération Francophone de Cancérologie Digestive (FFCD). They are also in part based on public data generated by 10X Genomics: https://www.10xgenomics.com/datasets, and by the TCGA Research Network: https://www.cancer.gov/tcga. We thank Daniel Gonzalez, Jean-Philippe Vert and Alberto Romagnoni for their support and insightful discussions.

## Contributors

BS, LH, AO, TD, RD, LG, JBS. and VDP. wrote the code, performed the experiments and analyzed the results. PLP provided the data (PETACC-8). DLC prepared the samples. BS, LH, AO, TD, LG, EP and ED wrote the manuscript with the assistance and feedback of all the other co-authors.

## Methods

### Data

#### PETACC8-Visium

The cohort used to train the model contained 48 slides of distinct Colon Adenocarcinoma patients with 10X Genomics Visium spatial transcriptomics from a subset of the PETACC 8 cohort [Taieb et al]^38^. Both Haematoxylin, Eosin & Saffron (HES)-stained slides’ images and Visium capture area images were available. On average, samples contained around 4,000 spots, and the expressions of up to 15,000 genes were measured in each spot. HES slides were scanned at a resolution of 0.5 micron per pixel.

24 new samples were made available after model development and were used as an external validation cohort. HES slides for these samples were scanned at a resolution of 0.9 micron per pixel.

#### Xenium samples

The Xenium samples we used were made publicly available by 10X genomics. For each sample, transcripts and cell coordinates, immunofluorescence (IF) image and H&E stained slide were available. A panel of 480 genes, specific to each sample, were sequenced with 10X Xenium technology.

#### TCGA dataset

This study made use of three publicly available dataset from The Cancer Genome Atlas Program (TCGA): Colon adenocarcinoma patients (TCGA COAD), Breast invasive carcinoma (TCGA-BRCA) and Stomach adenocarcinoma (TCGA-STAD). We selected samples from primary tumors only, for which diagnostic slides were available. Slides we used were digitized H&E-stained formalin-fixed, paraffin-embedded (FFPE) histology slides.

### Preprocessing of spatial transcriptomic data

The pre-processing of the spatial-transcriptome profiles were done using 10X Genomics Space Ranger software. Raw sequenced reads were aligned on a probe reference. Then we use our in-house matter detector to remove spots for which the histology image displayed a lack of matter.

For counts normalization, Bhuva *et al*.^39^ showed that library size is associated with tissue structure and that usual corrections like CPM or SCTransform could result in loss of information. As such we chose to use raw counts and employ a cosine loss function to correct for sample variations.

We selected 3 different lists of genes for prediction (see. supplementary materials). For benchmarking we used lists of the 100 and 5000 most varying genes over the training dataset computed with Moran’s I and aggregated with a geometric mean. For comparison with nuclei cell types annotations, we selected a curated list of 14 genes associated with the 14 cell types annotated.

### Preprocessing of histology slides

Histology slides are high dimensionality data, with up to 100 000 × 100 000 pixels for a single whole-slide image. For deep learning applications, it has thus become standard to divide the whole-slide image into hundreds to thousands of subparts referred to as tiles. In our work, we applied the same approach by extracting square images of size 224 × 224 pixels (approximately 112 × 112 μm) centered on each of the ∼5000 spots. Given that, in the 10x Genomics Visium technology, each spot measures 55 μm and that two spots are 100 μm apart, this enables a dense covering of the slide while capturing the image information associated with each sequenced spot. In addition to the Space Ranger matter detection, we also used an in-house deep learning matter detection model, trained internally on a dataset of manually annotated whole-slide images. This enabled a finer selection of tiles that contain matter and removed artifacts.

For each tile, a 768-dimension feature descriptive vector was extracted using an in-house pre-trained deep learning model^40^: a base vision-transformer model trained on TCGA-COAD using the IBOT Self-Supervised Learning framework. We used those tile-level representations for the supervised approach. For the weakly-supervised approach, we needed to obtain patch-level descriptive vectors. By design, a vision transformer separates an input image into several patches, in our case 196 patches of 16 × 16 pixels. During the self-supervised training, a class token is used to extract a tile-level representation, which is the one we used for the tile-level prediction task, but patch-level descriptors are also trained within that framework through the Masked Image Modeling loss. Thus, for the weakly supervised approach, we used those ones as the patch descriptive vectors.

Since the slides from the validation cohort were scanned at a lower resolution, tile images of size 112 × 112 pixels were resized to 224 × 224 before feature extraction.

### Modeling of Spatial Transcriptomic data

As outlined in previous work^41^, counts in spatial transcriptomic data should follow a Poisson law

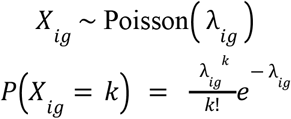

Here, *X*_*ig*_ is the measured count for gene *g* in spot *i*, while *λ*_*ig*_ is an unobserved latent variable that can depend both on underlying biology (for instance, the cell composition in a spot) and on technical effects such as inter-spot contamination. Since all spots are different, the distribution of counts for a given gene on a slide should behave as a mixture-of-Poisson. This is often modeled as a negative binomial distribution, which is a special case of mixture-of-Poisson where the parameter *λ* follows a Gamma law.

In practice, we observe that genes with little expression tend to follow a Poisson law over the slide, meaning that they show no spatial heterogeneity, while genes with larger counts are better modeled by a negative binomial distribution.

An important point of the Poisson distribution is that its mean μ and variance σ^2^ are equal, while negative binomial distribution is characterized by an overdispersion 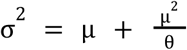.

Since the quantity we measure is not the biological ground truth, but rather a random variable with statistical fluctuations, we want to evaluate how much signal is contained in the data. A way to measure this is to evaluate the Pearson correlation between the unobserved latent variable *λ*_*ig*_ that contains the real signal, and the measured counts *X*_*ig*_. Based on the relation between *X*_*ig*_ and *λ*_*ig*_, one can show that this correlation is given by

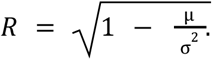

This formula is a way to measure how much the distribution of counts over the slide deviates from a pure Poisson distribution. Whenever *R* = 0, it means that *λ*_*ig*_ does not depend on *i*, meaning the distribution over the slide is a Poisson. In this case, the only variations observed are due to statistical noise and there is no spatial signal.

One can also show that, for a given spatial distribution of a gene, *R*_*max*_ is a growing function of the sequencing depth, which can be modeled as a factor scaling the distribution parameters *λ*_*ig*_.

This quantity, referred to as *R*_*max*_, is an estimate of the maximum Pearson correlation we could achieve with MISO, because a perfect oracle, knowing exactly the underlying variation factors, would reach this performance.

### Selection of genes for enrichment analysis

When evaluating performance of our models, for each (gene, sample) pair, we compared the measured Pearson correlation (average over test folds where the sample appeared) between the prediction and the labels to this theoretical upper bound. In almost every case, correlation fell below *R*_*max*_, with very few exceptions being observed exclusively for low values (*R*_*max*_ < 0.2), probably as a result of spurious correlations. In order to evaluate the ability of the model to predict biological features independently from the noise contained in the data, we computed the ratio *R*_*obs*_ /*R*_*max*_ After excluding pairs (gene, slide) for which there was too little signal (*R*_*max*_ < 0.2), we selected pairs that reached a ratio *R*_*obs*_ /*R*_*max*_, and ranked genes by the number of times they appeared in this list. 118 genes were consistently well predicted, with a ratio above 0.5 on 18 samples or mores. Those were selected for gene set enrichment analysis.

### Models for Spatial Transcriptomic prediction

The baseline architecture consists essentially of an MLP, that processes in parallel every spot, to predict gene expression. We considered a baseline with 2 layers and respectively 1024 and 512 hidden units per layer.

Self-attention in transformer networks is defined by a scaled dot-product attention. In our case, the feature vector representing a tile *x*_*i*_ is passed through three linear layers to generate three vectors named key (*K*_*i*_), query (*Q*_*i*_) and value (*V*_*i*_). The self-attention weight between two tiles *i* and *j* is defined as

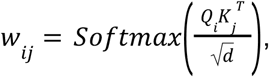

where *d* is the dimension of the key and query vectors. The output representation of the self-attention module is defined as

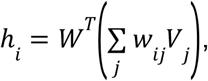

where *W* is a projection matrix. The full transformer block is defined by

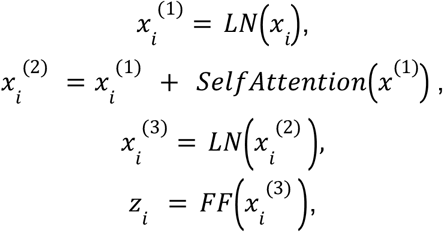

where *LN* denotes a Layer Norm operator^42^ and *FF* a feedforward neural network (a single linear layer in our case).

Local Attention Multiple Instance Learning was introduced in previous work by Reisenbüchler et al.^26^. It consists of a transformer-like architecture, in which self-attention is not computed for every pair of instances, but only for neighboring instances. This means that, in the previous equations, *w*_*ij*_ is zero except if tile *j* is one of the *k* nearest neighbors of tile *i*, based on euclidean distance in the space of tile coordinates.

Here, we considered an architecture with a first feature embedding layer with 1024 hidden units, followed by one transformer block with internal dimension 512 and 32 attention heads. Another densely connected layer was used to map the output of the transformer block to 1024, and a last linear layer with dimension equal to the number of genes was applied for gene expression prediction. The number of neighbors used in attention computation was optimized by cross-validation.

The weakly supervised approach follows the idea of a two-step prediction. For each pair of sequenced spot and tile image, we divided the 224 × 224 pixel image into 196 adjacent patches, of size 16 × 16 pixels (approximately 8 × 8 μm). We then modified the baseline architecture by making the MLP prediction at the patch level and added an aggregation mechanism (average pooling) during training. Then, the loss function was calculated on those aggregated predictions.

### Loss function

Raw spatial transcriptomic data present strong inter-sample variations, as the sequencing depth for a given gene may vary a lot across slides. Contrary to bulk and single-cell transcriptomic, so far, no consensus method has emerged yet to normalize this kind of data. As a consequence, training a model to reproduce raw counts by minimizing Mean Squared Error (MSE) might not be optimal, as the scale will be different from one sample to another. To overcome this issue while making minimal assumptions about the data, we chose instead to train the model directly to predict the relative expression of a given gene across spots from the same sample. We defined a loss function based on cosine similarity, that is invariant under any rescaling of the labels and predictions. During training, all spots from a given slide were processed in the same batch, the cosine similarity between predictions and labels was computed separately for each slide of the batch, and averaged over slide. To maximize cosine similarity by gradient descent, the loss function was defined as

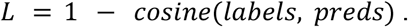

Experiments with the baseline architecture (MLP) demonstrated superior performance of our cosine similarity-based loss compared to MSE (Average Spearman correlation of 0.343 and 0.296 respectively, p-value < 10^−4^, Fig. S1).

### Distillation and super-resolution

When performing weakly supervised learning for super-resolution, the number of instances in a given slide was multiplied by 196 (the number of patches in a tile image), making it computationally challenging to go through the same training procedure. We overcame this by using a distillation technique. For a given list of genes, the LAMIL architecture (the teacher model) was first trained with cosine similarity-based loss, then the predictions of this architecture were used as labels for training the weakly supervised model (the student model) with MSE. The intuition was that the predictions of a model trained with cosine similarity-based loss would be rid of technical variation across samples. The metrics of the weakly-supervised model were still computed against the raw Visium counts. The comparison of spot-level performances showed the superiority of this approach against a naïve one in which raw counts were directly predicted by the weakly supervised model using MSE.

We found that knowledge transfer significantly improved the performance of our weakly supervised model with respect to *de novo* training, from a spearman correlation of 0.292 (0.049) to 0.373 (0.045) in internal validation, and 0.333 to 0.433 in external validation to predict the expression of the 100 genes with highest Moran’s I (table 3).

**Table 3:**
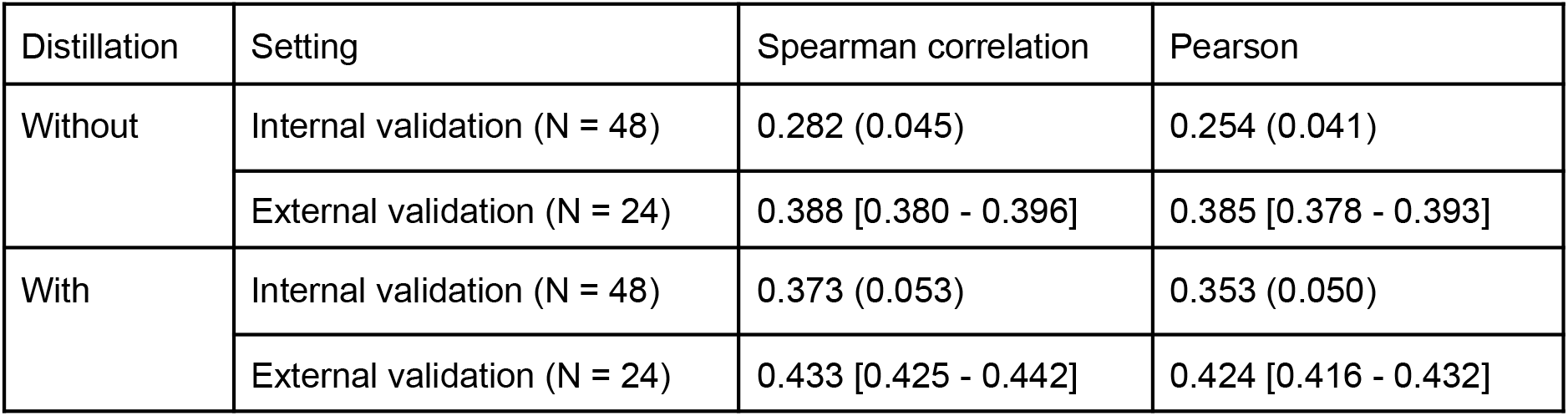
Spot-level performances of the weakly supervised models trained with or without knowledge distillation.

| Distillation | Setting | Spearman correlation | Pearson |
| --- | --- | --- | --- |
| Without | Internal validation (N = 48) | 0.282 (0.045) | 0.254 (0.041) |
|  | External validation (N = 24) | 0.388 [0.380 - 0.396] | 0.385 [0.378 - 0.393] |
| With | Internal validation (N = 48) | 0.373 (0.053) | 0.353 (0.050) |
|  | External validation (N = 24) | 0.433 [0.425 - 0.442] | 0.424 [0.416 - 0.432] |

**Table 4:** description of the cohorts used for transfer learning tasks.

| Cohort | Task | Number of patients | Distribution |
| --- | --- | --- | --- |
| TCGA-COAD | MSI | 408 | 73 MSI-H<br>335 MSI-L and MSS |
|  | CMS | 318 | 62 CMS1<br>113 CMS2<br>55 CMS3<br>88 CMS4 |
|  | OS | 431 | 97 observed events<br>334 censors |
|  | Bulk gene expression prediction | 431 |  |
| TCGA-STAD | MSI | 375 | 64 MSI-H<br>311 MSI-L and MSS |
|  | OS | 370 | 143 observed events<br>227 censors |
| TCGA-BRCA | Molecular subtype | 1007 | 537 Luminal A<br>215 Luminal B<br>79 HER2-enriched<br>176 Basal-like |
|  | HRD | 1003 | 198 HRD<br>805 HRP |
|  | BRCA1/BRCA2 | 1015 | 50 BRCA1/BRCA2 mutated<br>965 wild-type |
|  | OS | 1050 | 145 observed events<br>905 censors |
|  | Bulk gene expression prediction | 1048 |  |

### iStar

As the iStar method was designed to make predictions on either the training slide or a consecutive one only, a small adaptation of the method was needed to evaluate it in the same setting as MISO. The iStar model is trained to predict rescaled count values (using min-max scaling on a per-gene basis, computed over the training sample). During inference, the predicted counts are rescaled back to their correct original range. In our external inference scenario, we perform the min-max scaling using all the samples from the training set and use these min-max values to scale back the predictions at inference, without guarantees that this range is correct due to the significant inter-patient variability. This strategy prevents us from leaking inference-time information into the validation. All the other parameters (for data preprocessing and training) are the same as in the original implementation. Furthermore, the performances reported are averages from models trained on the same 25 random train/test splits of the data used for MISO.

### Annotation of nuclei

11640 cells, across 126 tiles from 38 slides of the TCGA-COAD cohort were internally annotated. This annotation process was done by 7 pathologists hired for the task with several redundancies to maximize agreements between experts. Each tile has a size of 448 × 448 pixels and was annotated at the highest resolution level available (0.25 µm per pixel). The annotated tiles were selected in order to maximize distinct cell type populations, as well as the presence of low-abundance populations such as eosinophils and neutrophils. The annotation process was the following: pathologists point the approximate center of each cell and register their identified cell type. Afterwards, the interactive nuclei segmentation framework NuClick^43^ is used to infer segmentation around each individual point. Cell types annotated were: Apoptotic Body, Cancer cell, Endothelial Cell, Eosinophils, Epithelial cell (non-cancerous), Fibroblasts, Macrophages, Red blood cell, Plasmocytes and Lymphocytes.

### Transfer learning

For transfer learning, MISO was trained on the whole development cohort (48 slides) and applied to TCGA-COAD. The prediction layer was removed, to extract a representation of size 1024 from each tile.

All models trained in cross-validation for downstream tasks had the same architecture that was not further optimized. Models were based on the Chowder architecture ^33^. Each model had 5 scoring channels, 50 copies of the models with different weight initialization were trained simultaneously and their predictions were averaged at test time, as this has been shown to increase the robustness of the results. On each fold, models were trained for 10 epochs.

Models were trained on the whole cohort with 5-fold cross-validation, repeated 20 times with different random seeds to ensure the reliability of the comparison. For each task, in order to evaluate each representation as a function of the dataset size, we also subsampled different numbers n of patients from the target domain cohort. For TCGA-COAD and TCGA-STAD, we considered numbers of patients in (25, 50, 75, 100, 150, 200, 300), while for TCGA-BRCA we considered n in (50, 100, 200, 300, 500). For each value of *n*, the dataset was sampled 20 times with different random seeds, and a 5-fold cross-validation was run on the subsampled dataset, to produce a total of 100 models per dataset size.

We trained models on four different kinds of tasks:

- Bulk gene expression prediction: we considered lists of genes associated with hallmarks of cancer (cell cycle, DNA repair, angiogenesis, hypoxia, immune response mediated by B cells, and adaptive immune response mediated by T cells)^44^, as well as marker genes of lymphocytes (*CD3D, CD3E, CD3G, CD247* for T cells, *CD19* and *MS4A1* for B cells).
- Molecular subtyping: prediction of Consensus Molecular Subtypes (CMS) in colorectal cancer^45^, and prediction of molecular subtypes of breast cancer (Luminal A, Luminal B, HER2-enriched and Basal)^46^.
- Genetic alterations: prediction of MicroSatellite Instability (MSI)^47^ in colon and gastric cancer, Homologous Repair Deficiency (HRD)^48^ and BRCA1/BRCA2 mutation^49^ in breast cancer.
- Survival: prediction of Overall Survival (OS).

The dataset size for each task is indicated in the table below.

### Statistics and p-values

To compare the correlations achieved by two models on the same gene and sample, we used a one-tailed t-test on Fisher z-transformed correlation coefficients^50^ (Hinkel et al, 1988). Aggregation of p-values over repeated train/test splits was done by computing the median p-value and multiplying by 2.

To compare the performances of models trained on top of iBOTViTBase representation or MISO representation, for every task and every training dataset size, we ran a corrected resampled T-test^51^ on the 20 × 5 metric values. This procedure accounts for the non-independence between the folds of the repeated cross-validation, by correcting the estimated variance, defined as

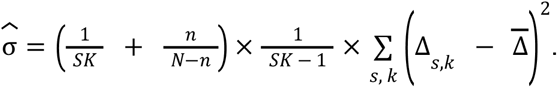

*S* is the number of repeats (here, *S* = 20), *K* the number of folds (*K* = 5), N the total size of the cohort, n the size of the test set, Δ_*s,k*_ the difference between the performances of the two models on repeat *s*, fold *k* and 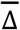 the average difference. The p-value is obtained by running a one-tailed t-test on the rescaled difference.

Combination of p-values over different tasks was performed with Stouffer’s method^52^.

## Bibliography

1. Wang, Z., Gerstein, M. & Snyder, M. RNA-Seq: a revolutionary tool for transcriptomics. Nat. Rev. Genet. 10, 57–63 (2009).

2. Zhao, H. et al. Whole transcriptome RNA-seq analysis: tumorigenesis and metastasis of melanoma. Gene 548, 234–243 (2014).

3. Zhang, E. et al. A novel long noncoding RNA HOXC-AS3 mediates tumorigenesis of gastric cancer by binding to YBX1. Genome Biol. 19, 154 (2018).

4. Wang, J., Dean, D. C., Hornicek, F. J., Shi, H. & Duan, Z. RNA sequencing (RNA-Seq) and its application in ovarian cancer. Gynecol. Oncol. 152, 194–201 (2019).

5. Eswaran, J. et al. Transcriptomic landscape of breast cancers through mRNA sequencing. Sci. Rep. 2, 264 (2012).

6. Cui, A. et al. Dictionary of immune responses to cytokines at single-cell resolution. Nature 625, 377–384 (2024).

7. Xie, Y. et al. A global database for modeling tumor-immune cell communication. Sci. Data 10, 444 (2023).

8. Sheng, J. et al. Topological analysis of hepatocellular carcinoma tumour microenvironment based on imaging mass cytometry reveals cellular neighbourhood regulated reversely by macrophages with different ontogeny. Gut 71, 1176–1191 (2022).

9. Risom, T. et al. Transition to invasive breast cancer is associated with progressive changes in the structure and composition of tumor stroma. Cell 185, 299-310.e18 (2022).

10. Zidane, M. et al. A review on deep learning applications in highly multiplexed tissue imaging data analysis. Front. Bioinforma. 3, (2023).

11. Coudray, N. et al. Classification and mutation prediction from non–small cell lung cancer histopathology images using deep learning. Nat. Med. 24, 1559–1567 (2018).

12. Kather, J. N. et al. Deep learning can predict microsatellite instability directly from histology in gastrointestinal cancer. Nat. Med. 25, 1054–1056 (2019).

13. Saillard, C. et al. Pacpaint: a histology-based deep learning model uncovers the extensive intratumor molecular heterogeneity of pancreatic adenocarcinoma. Nat. Commun. 14, 3459 (2023).

14. Schmauch, B. et al. A deep learning model to predict RNA-Seq expression of tumours from whole slide images. Nat. Commun. 11, 3877 (2020).

15. Courtiol, P. et al. Deep learning-based classification of mesothelioma improves prediction of patient outcome. Nat. Med. 25, 1519–1525 (2019).

16. He, B. et al. Integrating spatial gene expression and breast tumour morphology via deep learning. Nat. Biomed. Eng. 4, 827–834 (2020).

17. Monjo, T., Koido, M., Nagasawa, S., Suzuki, Y. & Kamatani, Y. Efficient prediction of a spatial transcriptomics profile better characterizes breast cancer tissue sections without costly experimentation. Sci. Rep. 12, 4133 (2022).

18. Mejia, G., Cárdenas, P., Ruiz, D., Castillo, A. & Arbeláez, P. SEPAL: Spatial Gene Expression Prediction from Local Graphs. Preprint at http://arxiv.org/abs/2309.01036 (2024).

19. Zeng, Y. et al. Spatial transcriptomics prediction from histology jointly through Transformer and graph neural networks. Brief. Bioinform. 23, bbac297 (2022).

20. Jia, Y., Liu, J., Chen, L., Zhao, T. & Wang, Y. THItoGene: a deep learning method for predicting spatial transcriptomics from histological images. Brief. Bioinform. 25, bbad464 (2023).

21. Xiao, X., Kong, Y., Li, R., Wang, Z. & Lu, H. Transformer with convolution and graph-node co-embedding: An accurate and interpretable vision backbone for predicting gene expressions from local histopathological image. Med. Image Anal. 91, 103040 (2024).

22. Ståhl, P. L. et al. Visualization and analysis of gene expression in tissue sections by spatial transcriptomics. Science 353, 78–82 (2016).

23. Andersson, A. et al. Spatial deconvolution of HER2-positive breast cancer delineates tumor-associated cell type interactions. Nat. Commun. 12, 6012 (2021).

24. Ji, A. L. et al. Multimodal Analysis of Composition and Spatial Architecture in Human Squamous Cell Carcinoma. Cell 182, 497–514.e22 (2020).

25. Jaume, G. et al. HEST-1k: A Dataset for Spatial Transcriptomics and Histology Image Analysis. Preprint at http://arxiv.org/abs/2406.16192 (2024).

26. Reisenbüchler, D., Wagner, S. J., Boxberg, M. & Peng, T. Local Attention Graph-Based Transformer for Multi-target Genetic Alteration Prediction. in Medical Image Computing and Computer Assisted Intervention – MICCAI 2022 (eds. Wang, L., Dou, Q., Fletcher, P. T., Speidel, S. & Li, S.) vol. 13432 377–386 (Springer Nature Switzerland, Cham, 2022).

27. Hafemeister, C. & Satija, R. Normalization and variance stabilization of single-cell RNA-seq data using regularized negative binomial regression. Genome Biol. 20, 296 (2019).

28. Satoh, K. et al. Global metabolic reprogramming of colorectal cancer occurs at adenoma stage and is induced by MYC. Proc. Natl. Acad. Sci. 114, (2017).

29. Zhao, E. et al. Spatial transcriptomics at subspot resolution with BayesSpace. Nat. Biotechnol. 39, 1375–1384 (2021).

30. Bergenstråhle, L. et al. Super-resolved spatial transcriptomics by deep data fusion. Nat. Biotechnol. 40, 476–479 (2022).

31. Zhang, D. et al. Inferring super-resolution tissue architecture by integrating spatial transcriptomics with histology. Nat. Biotechnol. (2024) doi:10.1038/s41587-023-02019-9.

32. Hinton, G., Vinyals, O. & Dean, J. Distilling the Knowledge in a Neural Network. Preprint at http://arxiv.org/abs/1503.02531 (2015).

33. Courtiol, P., Tramel, E. W., Sanselme, M. & Wainrib, G. Classification and Disease Localization in Histopathology Using Only Global Labels: A Weakly-Supervised Approach. Preprint at http://arxiv.org/abs/1802.02212 (2020).

34. Lengrand, J. et al. Pharmacological targeting of netrin-1 inhibits EMT in cancer. Nature 620, 402–408 (2023).

35. Cassier, P. A. et al. Netrin-1 blockade inhibits tumour growth and EMT features in endometrial cancer. Nature 620, 409–416 (2023).

36. Madissoon, E. et al. A spatially resolved atlas of the human lung characterizes a gland-associated immune niche. Nat. Genet. 55, 66–77 (2023).

37. Lyubetskaya, A. et al. Assessment of spatial transcriptomics for oncology discovery. Cell Rep. Methods 2, 100340 (2022).

38. Taieb, J. et al. Oxaliplatin, fluorouracil, and leucovorin with or without cetuximab in patients with resected stage III colon cancer (PETACC-8): an open-label, randomised phase 3 trial. Lancet Oncol. 15, 862–873 (2014).

39. Bhuva, D. D. et al. Library size confounds biology in spatial transcriptomics data. Genome Biol. 25, 99 (2024).

40. Filiot, A. et al. Scaling Self-Supervised Learning for Histopathology with Masked Image Modeling. 2023.07.21.23292757 Preprint at 10.1101/2023.07.21.23292757 (2023).

41. Zhao, P., Zhu, J., Ma, Y. & Zhou, X. Modeling zero inflation is not necessary for spatial transcriptomics. Genome Biol. 23, 118 (2022).

42. Ba, J. L., Kiros, J. R. & Hinton, G. E. Layer Normalization. Preprint at http://arxiv.org/abs/1607.06450 (2016).

43. Koohbanani, N. A., Jahanifar, M., Tajadin, N. Z. & Rajpoot, N. NuClick: A Deep Learning Framework for Interactive Segmentation of Microscopy Images. Preprint at http://arxiv.org/abs/2005.14511 (2020).

44. Hanahan, D. & Weinberg, R. A. Hallmarks of cancer: the next generation. Cell 144, 646–674 (2011).

45. Thanki, K. et al. Consensus Molecular Subtypes of Colorectal Cancer and their Clinical Implications. Int. Biol. Biomed. J. 3, 105–111 (2017).

46. Eliyatkin, N., Yalcin, E., Zengel, B., Aktaş, S. & Vardar, E. Molecular Classification of Breast Carcinoma: From Traditional, Old-Fashioned Way to A New Age, and A New Way. J. Breast Health 11, 59–66 (2015).

47. Boland, C. R. & Goel, A. Microsatellite Instability in Colorectal Cancer. Gastroenterology 138, 2073-2087.e3 (2010).

48. Stewart, M. D. et al. Homologous Recombination Deficiency: Concepts, Definitions, and Assays. The Oncologist 27, 167–174 (2022).

49. Baretta, Z., Mocellin, S., Goldin, E., Olopade, O. I. & Huo, D. Effect of BRCA germline mutations on breast cancer prognosis: A systematic review and meta-analysis. Medicine (Baltimore) 95, e4975 (2016).

50. Witz, K., Hinkle, D. E., Wiersma, W. & Jurs, S. G. Applied Statistics for the Behavioral Sciences. J. Educ. Stat. 15, 84 (1990).

51. Nadeau, C. & Bengio, Y. Inference for the Generalization Error. Mach. Learn. 52, 239–281 (2003).

52. Stouffer, S. A., Suchman, E. A., Devinney, L. C., Star, S. A. & Williams Jr., R. M. The American Soldier: Adjustment during Army Life. (Studies in Social Psychology in World War II), Vol. 1. xii, 599 (Princeton Univ. Press, Oxford, England, 1949).

