## Supplementary Material for "A deep learning-based multiscale integration of spatial omics with tumor morphology"

### Supplementary Table 1

| | gene | Spearman | Pearson |
|---|---|---|---|
| Genes with Spearman correlation above 0.5 | *EPCAM* | 0.721 | 0.631 |
| | *GPX2* | 0.717 | 0.599 |
| | *ERBB3* | 0.672 | 0.561 |
| | *GPRC5A* | 0.663 | 0.522 |
| | *RNF43* | 0.640 | 0.555 |
| | *CDX2* | 0.623 | 0.492 |
| | *MET* | 0.613 | 0.488 |
| | *CEACAM1* | 0.604 | 0.452 |
| | *SOX9* | 0.571 | 0.465 |
| | *ERBB2* | 0.566 | 0.483 |
| | *TMC5* | 0.544 | 0.459 |
| | *TP53* | 0.538 | 0.445 |
| | *CTNNB1* | 0.535 | 0.451 |
| | *ACTB* | 0.526 | 0.424 |
| | *CD44* | 0.522 | 0.449 |
| | *PPARG* | 0.516 | 0.400 |
| | *TFF3* | 0.516 | 0.348 |
| | *IFITM3* | 0.514 | 0.434 |
| | *CD47* | 0.513 | 0.459 |
| | *SEMA3C* | 0.508 | 0.424 |
| | *SDC1* | 0.507 | 0.435 |
| T cell and B cell marker genes | *CD3D* | 0.160 | 0.203 |
| | *CD3E* | 0.208 | 0.263 |
| | *CD247* | 0.105 | 0.122 |
| | *CD19* | 0.065 | 0.120 |

Table S1: Performance of patch-level (8μm x 8μm) predictions for the best predicted genes and B and T cell markers on the Xenium Colorectal Cancer sample.

### Supplementary Table 2

| **targeted cells** | **genes Xenium panel** | **genes model** | **Patch size (μm²)** | **Spearman** | **Pearson** |
|---|---|---|---|---|---|
| T cells | *CD3D* *CD3E* *CD247* | *CD3D* *CD3E* *CD3G* *CD247* | 8x8 | 0.237 | 0.296 |
| | | | 16x16 | 0.396 | 0.411 |
| | | | 32x32 | 0.543 | 0.507 |
| | | | 56x56 | 0.599 | 0.537 |
| B cells | *CD19* *MS4A1* | *CD19* | 8x8 | 0.100 | 0.198 |
| | | | 16x16 | 0.165 | 0.298 |
| | | | 32x32 | 0.226 | 0.365 |
| | | | 56x56 | 0.305 | 0.367 |
| T and B cells | *CD3D* *CD3E* *CD247* *CD19* *MS4A1* | *CD3D* *CD3E* *CD3G* *CD247* *CD19* | 8x8 | 0.260 | 0.360 |
| | | | 16x16 | 0.416 | 0.468 |
| | | | 32x32 | 0.557 | 0.537 |
| | | | 56x56 | 0.618 | 0.569 |

Table S2: Patch-level performance for the prediction of gene expression summed over T cell and B cell markers, with varying patch size, Xenium colorectal cancer sample.

### Supplementary Table 3

| | | 6μm x 6μm patches | | 12μm x 12μm patches | |
|---|---|---|---|---|---|
| | **gene** | **Spearman** | **Pearson** | **Spearman** | **Pearson** |
| Genes with Spearman correlation above 0.4 with 6μm x 6μm patches or above 0.5 with 12μm x 12μm patches | *EPCAM* | 0.501 | 0.495 | 0.562 | 0.621 |
| | *KRT8* | 0.498 | 0.458 | 0.622 | 0.639 |
| | *MLPH* | 0.468 | 0.510 | 0.576 | 0.634 |
| | *CD9* | 0.438 | 0.441 | 0.570 | 0.606 |
| | *SCD* | 0.430 | 0.419 | 0.496 | 0.529 |
| | *ERBB2* | 0.409 | 0.469 | 0.499 | 0.585 |
| | *CDH1* | 0.407 | 0.385 | 0.537 | 0.552 |
| | *FASN* | 0.407 | 0.407 | 0.523 | 0.562 |
| | *MYO5B* | 0.335 | 0.329 | 0.546 | 0.555 |
| | *DSP* | 0.355 | 0.338 | 0.542 | 0.543 |
| | *ELF3* | 0.306 | 0.331 | 0.516 | 0.546 |
| T cell and B cell marker genes | *CD3D* | 0.123 | 0.185 | 0.210 | 0.267 |
| | *CD3E* | 0.167 | 0.240 | 0.246 | 0.300 |
| | *CD3G* | 0.113 | 0.166 | 0.200 | 0.243 |
| | *CD247* | 0.118 | 0.177 | 0.201 | 0.243 |
| | *CD19* | 0.061 | 0.111 | 0.118 | 0.186 |

Table S3: Performance of patch-level predictions for the best predicted genes and B and T cell markers on the Xenium sample of breast carcinoma, with patches of size 6μm x 6μm (highest resolution) or 12μm x 12μm.

### Supplementary Table 4

| targeted cells | genes Xenium panel | genes model | Patch size (µm) | Spearman | Pearson |
|---|---|---|---|---|---|
| T cells | *CD3D*<br>*CD3E*<br>*CD3G*<br>*CD247* | *CD3D*<br>*CD3E*<br>*CD3G*<br>*CD247* | 6 | 0.1773 | 0.2970 |
| | | | 12 | 0.2437 | 0.3383 |
| | | | 23 | 0.2916 | 0.3413 |
| | | | 40 | 0.3463 | 0.3598 |
| B cells | *CD19*<br>*MS4A1* | *CD19* | 6 | 0.1062 | 0.1935 |
| | | | 12 | 0.1801 | 0.2863 |
| | | | 23 | 0.2772 | 0.3916 |
| | | | 40 | 0.3732 | 0.4576 |
| T and B cells | *CD3D*<br>*CD3E*<br>*CD3G*<br>*CD247*<br>*CD19*<br>*MS4A1* | *CD3D*<br>*CD3E*<br>*CD3G*<br>*CD247*<br>*CD19* | 6 | 0.2080 | 0.3661 |
| | | | 12 | 0.2901 | 0.4226 |
| | | | 23 | 0.3610 | 0.4375 |
| | | | 40 | 0.4316 | 0.4702 |

Table S4: Patch-level performance for the prediction of gene expression summed over T cell and B cell markers, Xenium breast cancer sample.

### Supplementary Table 5

| Genes | | Colorectal cancer Xenium sample (8 µm patch) | | | Breast cancer Xenium sample (6 µm patch) | |
|---|---|---|---|---|---|---|
| | | Spearman | Pearson | | Spearman | Pearson |
| Top 5 predicted genes for iStar | *ERBB3* | 0.501 | 0.446 | *KRT8* | 0.480 | 0.450 |
| | *GPX2* | 0.446 | 0.421 | *SCD* | 0.472 | 0.473 |
| | *EPCAM* | 0.429 | 0.323 | *MLPH* | 0.471 | 0.455 |
| | *SOX9* | 0.420 | 0.357 | *EPCAM* | 0.448 | 0.416 |
| | *TMC5* | 0.417 | 0.371 | *FASN* | 0.441 | 0.461 |
| Avg. - 146 (resp. 105) genes | | 0.170 | 0.142 | | 0.139 | 0.136 |
| T and B cells markers (sum of expressions) | | 0.226 | 0.262 | | 0.091 | 0.107 |

Table S5: Patch-level performance of iStar on the Xenium samples. For B and T cell markers (*CD3*, *CD19*), we report the performance achieved to predict the sum of the expressions.

### Supplementary Figure 1

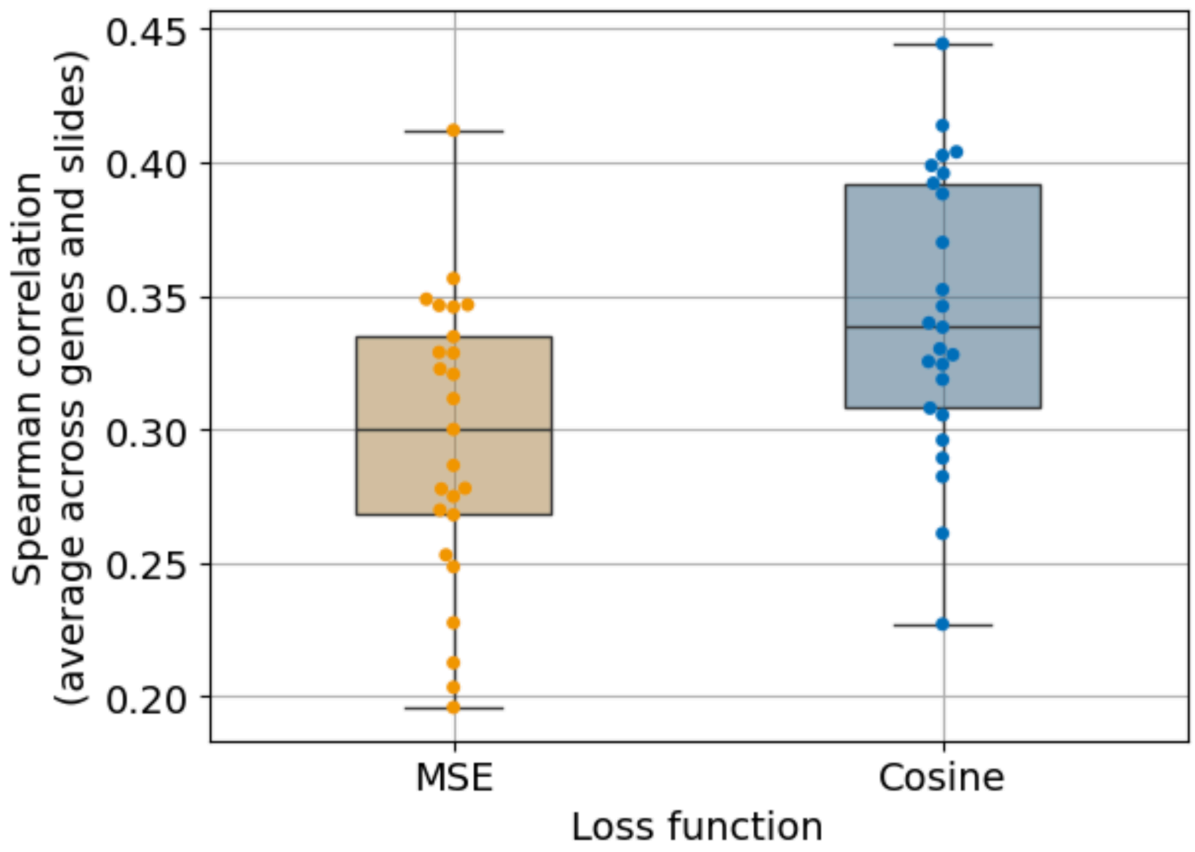


Figure S1: Performances (Spearman correlation) of the baseline architecture trained with Mean Squared Error (MSE) or with the loss based on cosine similarity.

### Supplementary Figure 2

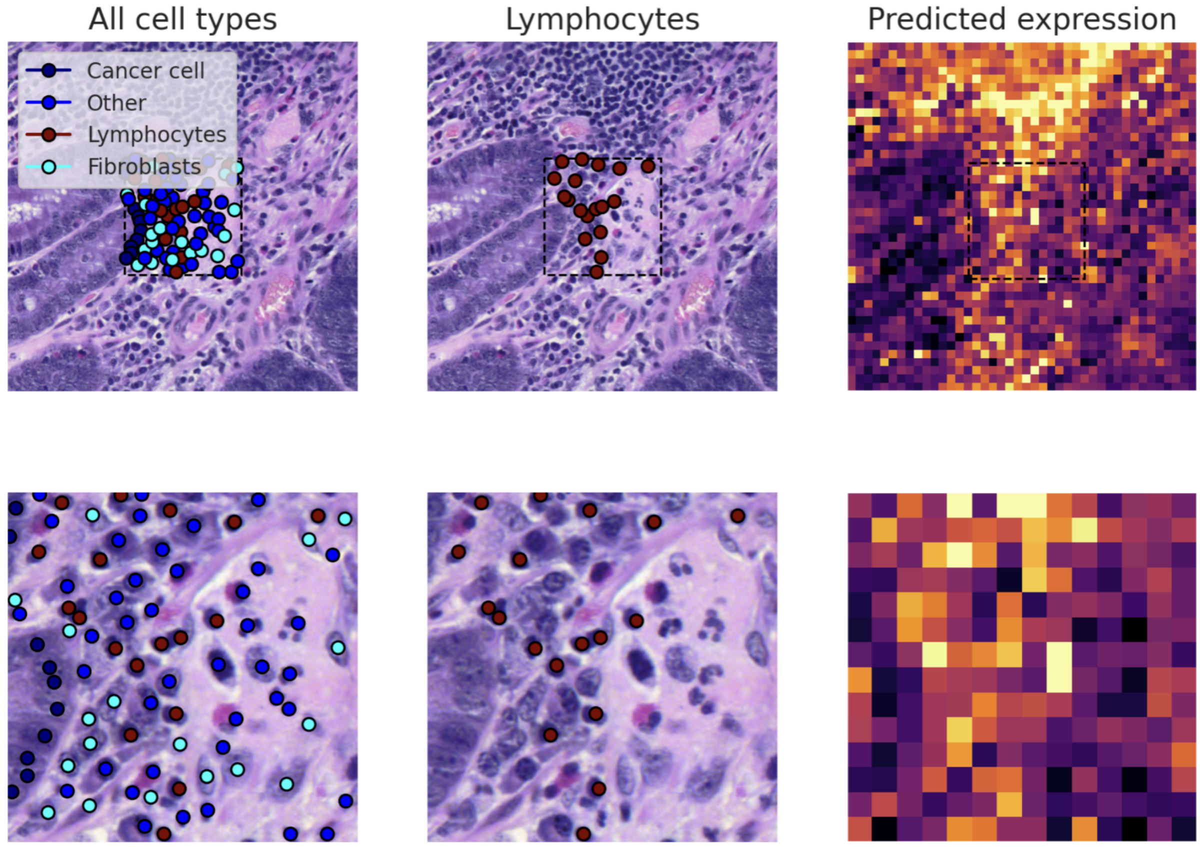


**Figure S3 - Predicted expression of *CD3*/*CD19* markers against cell type annotations**
Example tile annotated with lymphocytes and other cell types with (top) and without (bottom) its context, and predictions of the model for the *CD3D*, *CD3E*, *CD3G*, *CD247* and *CD19* genes (sum over genes) on patches of size 8µm x 8µm (left columns).

### Supplementary Figure 3

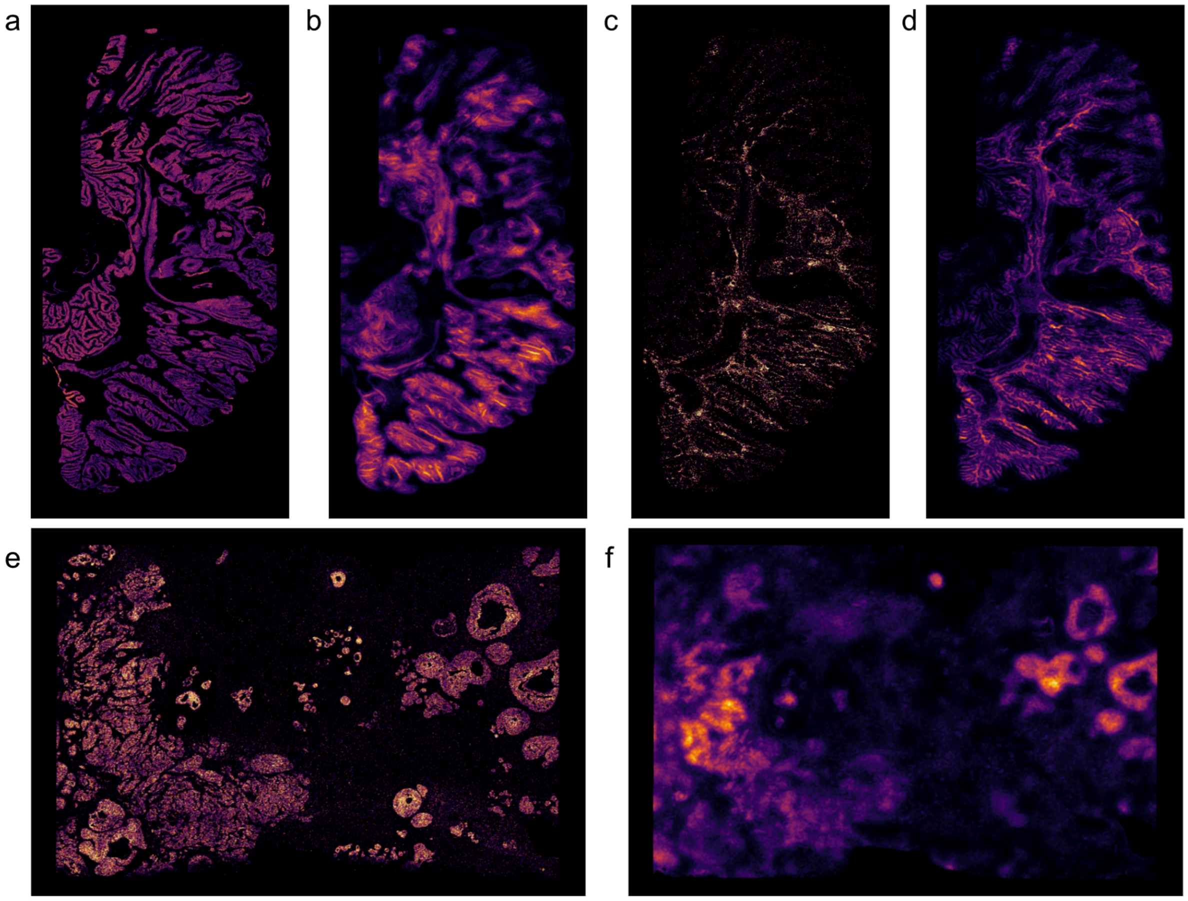


**Figure S3 - Validation of iStar on the Xenium samples.**

**a.** Ground truth expression of *EPCAM* on the colorectal cancer 10X Xenium sample. **b.** Predicted expression of *EPCAM* by the iStar method. **c.** Ground truth expression of T cell and B cell markers (average expression of *CD3D*, *CD3E*, *CD247*, *CD19 and MS4A1*) on the colorectal cancer 10X Xenium sample. **d.** Predicted expression of T cell and B cell markers (average expression of *CD3D*, *CD3E*, *CD3G*, *CD247* and *CD19*) by the iStar super-resolution method. **e.** Ground truth expression of *CDH1* on the breast cancer 10X Xenium sample. **f.** Prediction of the expression of *CDH1* (Spearman: 0.371, Pearson: 0.356) by the iStar method.
